# Isoform-selective NADPH oxidase inhibitor panel for pharmacological target validation

**DOI:** 10.1101/382226

**Authors:** V.T. Dao, Mahmoud H. Elbatreek, S. Altenhöfer, Ana I. Casas, M.P. Pachado, C.T. Neullens, U. Knaus, H.H.H.W. Schmidt

## Abstract

Unphysiological reactive oxygen species (ROS) formation is considered an important pathomechanism for several disease phenotypes with high unmet medical need. Therapeutically, antioxidants have failed multiple times. Instead, focusing on only disease-relevant, enzymatic sources of ROS appears to be a more promising and highly validated approach. Here the family of five NADPH oxidases (NOX) stands out as drug targets. Validation has been restricted, however, mainly to genetically modified rodents and is lacking in other species including human. It is thus unclear whether the different NOX isoforms are sufficiently distinct to allow selective pharmacological modulation. Here we show for five of the most advanced NOX inhibitors that indeed isoform selectivity can be achieved. NOX1 was most potently (IC_50_) targeted by ML171 (0.1 μM); NOX2, by VAS2870 (0.7 μM); NOX4, by M13 (0.01 μM) and NOX5, by ML090 (0.01 μM). Conditions need to be carefully controlled though as previously unrecognized non-specific antioxidant and assay artefacts may limit the interpretation of data and this included, surprisingly, one of the most advanced NOX inhibitors, GKT136901. As proof-of-principle that now also pharmacological and non-rodent target validation of different NOX isoforms is possible, we used a human blood-brain barrier model and NOX inhibitor panel at IC_50_ concentrations. The protective efficacy pattern of this panel confirmed the predominant role of NOX4 in stroke from previous genetic models. Our findings strongly encourage further lead optimization efforts for isoform-selective NOX inhibitors and clinical development and provide an experimental alternative when genetic validation of a NOX isoform is not an option.

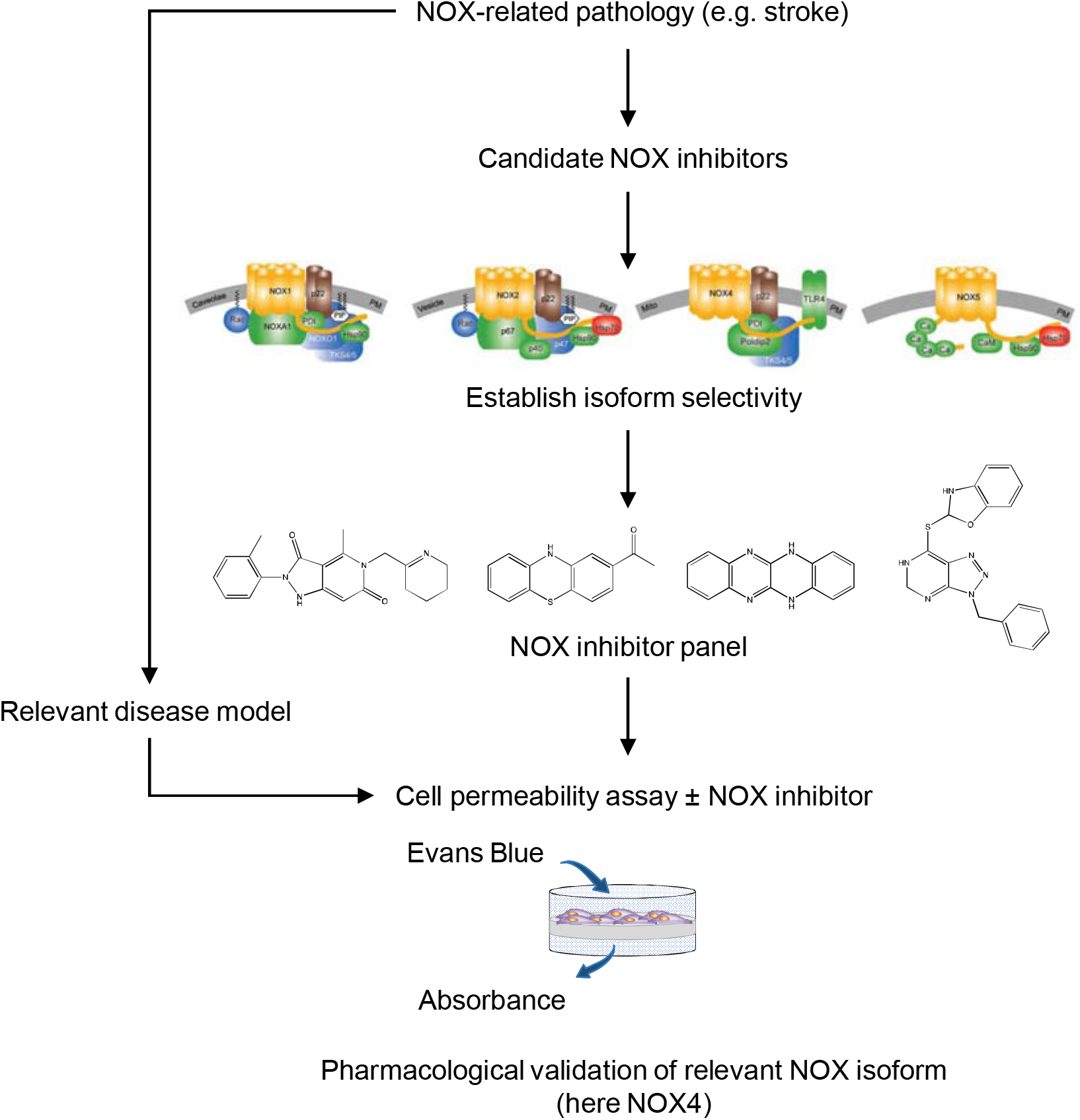
Graphical abstract.

## Introduction

Reactive oxygen species (ROS) are considered an important pathomechanism for several common diseases with high unmet medical need including cardiovascular and neurodegenerative as well as cancer. Yet, clinical translation of this hypothesis into therapeutic application by use of antioxidants that scavenge ROS has consistently failed [1], in some cases even increasing mortality [2–4]. This paradox was initially explained by these compounds being underdosed, thereby not reaching efficacy. It is now understood, however, that ROS are not only harmful metabolic by products, but also serve important protective, metabolic and signaling functions, such as the regulation of cell proliferation, differentiation, migration and survival, innate immune response, vascular tone, neuronal signaling as well as inflammation [5–8]. Anti-oxidants are likely to simultaneously interfere with both qualities of ROS, the physiological and pathophysiological ones with overall neutral or even deleterious outcomes. Thus, ROS should not be modulated in a systemic manner, but rather by identifying for each disease condition the relevant enzymatic ROS source and inhibit this source by selective pharmacological inhibitors [9].

Several enzymatic sources of ROS exist [10, 11]. NADPH oxidases (NOX), however, stand out as they are the only known dedicated, evolutionary conserved ROS-forming enzyme family. All other enzymes produce ROS as a byproduct or due to being damaged. Disease relevance has been suggested for NOX1, contributing to diabetic atherosclerosis [12] and retinopathy [13]; NOX2, neurodegeneration [14]; NOX4, stroke [15], diabetic nephropathy [16] and neuropathic pain [17], but protection against diabetic atherosclerosis [18, 19]; and NOX5, stroke [20], diabetic nephropathy [21, 22], hypertension [23] and coronary artery disease [24]. Target validation in these cases, however, has been done primarily by gene knock-out (as reviewed in [25]) and, in the case of NOX5 [21], by knock-in technology limiting most data sets to mice and, in the case of NOX4, in addition to rat [15].

For target validation in other species and eventual clinical translation, NOX-specific and ideally isoform-selective NOX inhibitors are desirable. It is unclear, however, whether this is or will be achievable given the fact that all five relevant NOX isoforms (NOX1, NOX2, NOX3, NOX4 and NOX5) are similarly structured transmembrane proteins containing highly conserved catalytic heme, FAD and NADPH binding sites. Some degree of variability derives from more (NOX1 and NOX2) or less (NOX4 and NOX5) binding partners or a unique regulation by calcium binding (NOX5) [26]. Furthermore, for each compound two potential sources of error need to be considered: (a) direct scavenging of ROS [27, 28] in instead of or in addition to NOX inhibition; and (b) interference with the ROS detection probes [29, 30]. Finally, from an efficacy and safety point of view whole cell-based assays provide additional information on membrane permeability and cytotoxicity.

Here, we examine some of the best characterized and/or most widely used NOX inhibitors, VAS2870, ML171, GKT136901, M13 and ML09 in a whole cell assay approach by analyzing isoform selectivity, possible assay interference and direct ROS-scavenging capacity. Finally, we suggest a NOX inhibitor/NOX isoform panel, which we then validate for usability in pharmacological target validation using human brain microvascular endothelial cell model of ischemia-induced hyperpermeability where NOX4 had previously been clearly validated as the key NOX isoform [15, 31].

## Material and Methods

### Chemicals and reagents

Dulbecco’s modified Eagle medium with GLUTAMAX (DMEM), RPMI-1640 medium with L-Glutamine and Hanks’ buffered salt solution (HBSS) were purchased from GIBCO/Life Technologies. Fetal bovine serum (FBS), 2-Acetylphenothiazine (ML171), 5,12-Dihydroquinoxalino(2,3-B)quinoxaline (ML090), phorbol myristate acetate (PMA), calcein, diphenyleneiodonium chloride (DPI), dextran sulfate, dimethyl sulfoxide (DMSO), Luminol, sodium salt, superoxide dismutase (SOD), G418, ionomycin calcium salt ready to made solution and Penicillin/Streptomycin solution were purchased from Sigma–Aldrich. 3-Benzyl-7-(2-benzoxazolyl) thio-1,2,3-triazolo [4,5-di] pyrimidine (VAS2870) was provided by Vasopharm; the pyrazolopyridine derivative GKT 136901, by GenKyoTex; M13, by Glucox Biotech. Amplex Red and horseradish peroxidase (HRP) were purchased from Invitrogen, CA, USA. The FuGENE6 transfection reagent was purchased from Promega.

### Cell culture and transfections

HEK293 cells were transfected with human NOXO1 (Gene ID: 124056), NOXA1 (Gene ID: 10811) and NOX1 (Gene ID: 27035) to measure NOX1 activity or individually with human NOX4 (Gene ID: 50507) or NOX5 (Gene ID: 79400) plasmids to measure NOX4 or NOX5 activities, respectively. All plasmids were confirmed by sequencing. Briefly, HEK293 cells were transfected with pcDNA control plasmid (vector control), NOX4 or NOX5 or triple transfected with NOX1, NOXO1 and NOXA1 using FuGENE6 transfection reagent (Promega) followed by ROS measurement after 48h. To measure NOX2 activity, as described previously [32], O_2_^•-^ was measured in HL-60 cells (human, promyeloblast, ATCC-No. CCL 240) that were cultured in RPMI-1640 medium with 5% FCS (JRH Biosciences), penicillin (100 U•mL-1), streptomycin (100 μg•mL-1) and glutamine (2 mM). Cell suspensions (26 × 10^6^ cells•mL-1) were incubated with 1.25% DMSO per 75cm^2^ TCF for 7 days to induce differentiation into granulocyte-like cells that were then centrifuged at 300xg, washed with HBSS and re-suspended in HBSS to the final seeding density needed for the experiment (5 × 10^5^ cells/well in 96 well plate).

### RNA Extraction and cDNA Synthesis

RNA was extracted using RNeasy^®^ Micro Kit (Qiagen) according to the manufacturer’s protocol. cDNA was synthesized from 1 μg total RNA in 20 μl reactions using High Capacity cDNA Reverse Transcription Kit (Thermo Fisher Scientific). After synthesis, the cDNA was stored at–20°C.

### Quantitative Real-Time PCR

RT-qPCR was performed on CFX96^™^ Real-Time PCR Detection System (Bio-Rad). All reactions were performed in triplicates in a total volume of 20 μl each using TaqMan^®^ Universal PCR Master Mix (Applied Biosystems-Life Technologies) according to manufacturer’s instructions. 3 μl cDNA was used as template and pre-designed TaqMan^®^ primers of β-actin, Nox1, Nox2, Nox4 and Nox5 were used. The specific assay ID for the primers used are shown in supplementary Table S1. The standard PCR conditions were as follows: 10 min at 95 °C, followed by 15 s at 95 °C and 1 min at 60 °C, 59 repeats. The amount of mRNA was normalized to the measured expression of β-actin mRNA.

### ROS measurement in intact cells

#### a. Luminol assay

To measure NOX1 activity, superoxide from vector-transfected or NOX1-transfected HEK293 cells was measured by luminol-enhanced chemiluminescence in white plates. For this assay HEK293 cells were transfected with NOX1, NOXO1 and NOXA1. After 48 hrs, cells were removed from the flasks by trypsinization and were re-suspended in PBS. 50 μl cell suspension consisting of 100,000 cells was added in triplicate to a 96-well plate in KRPG buffer and incubated at 37°C for 60 min. After 1 hr of incubation, 50 μl reaction buffer containing 6.4 U/ml HRP and 0.4 mM luminol in KRPG buffer was added. Cells were stimulated by addition of 0.5 μM Phorbol 12-myristate 13-acetate (PMA; PKC activator). ROS generation was detected by monitoring relative light units (RLU) with a Wallac luminometer Victor2 at 37°C for 60 min. Superoxide dismutase was added as control before PMA stimulation in selected wells.

#### b. Amplex Red assay

To measure the activity of NOX4 and NOX5, H_2_O_2_ production was measured in HEK293 cells using Amplex Red fluorescence. For the measurement, vector-transfected HEK293 cells or HEK293 cells transfected with NOX4 or NOX5 were trypsinized, counted, washed and re-suspended in Krebs-Ringer-Phosphate Glucose buffer (KRPG). NOX inhibitors dissolved in DMSO or solvent control (0.5% DMSO) were added to respective wells of a black 96-well plate and 50 μl of a reaction mixture containing 0.1 U·mL-1 HRP and 50 μM Amplex Red was added to each well according to the kit manual and incubated at 37°C for 10 min. Thereafter, 100,000 of respective HEK293 cells or solvent control in 50 μl of KPRG-buffer were added to each well. Amplex Red fluorescence was measured immediately at excitation (530-560 nm) and emission ~ 590 nm at 37°C for 60 min. HEK293 cells expressing NOX5 were activated by the Ca^2^+ ionophore ionomycin (40μM).

#### c. Cytochrome C assay

To measure NOX2 activity, superoxide production in HL-60 cells was measured by cytochrome C reduction as described previously [32]. Briefly, cytochrome C (100 μM) was added to the DMSO-differentiated HL-60 cell suspension. After adding 5 × 10^5^ cells to a 96-well plate and addition of inhibitors, a basal reading was performed at 540 nm (isosbestic point of cytochrome C) and 550 nm (SpectraMax 340; Molecular Devices, Sunnyvale, CA, USA). Subsequently, the oxidative burst was initiated by the addition of 100 nM PMA. After incubation for 60 min. at 37°C the absorbance was measured. Signals were normalized to the basal readings. In addition, superoxide dismutase was added as control before PMA stimulation in selected wells.

### Superoxide measurement in cell-free assays

Xanthine (X; final concentration: 50 μm) and cytochrome C (final concentration: 100 μm) were dissolved in HBSS, and 100 μL aliquots of this solution were transferred to individual wells of a 96-well plate. After addition of the oxidase inhibitors, the mixtures were allowed to equilibrate for 20 min. The reaction was started by the addition of 100 μL xanthine oxidase (XO) (final concentration: 5 mU·mL-1), and absorbance at 540 and 550 nm was recorded 10 min. after the start of the reaction. Superoxide production was calculated by normalization to the signals obtained at 540 nm. Similar experiments were performed with 0.4 mM luminol instead of cytochrome C. After 20 min. equilibration, the reaction was started with XOD (1 mU·mL-1), and chemiluminescence was subsequently recorded for 20 min. in a Fluoroscan FL microplate reader. Signals were calculated as AUC and normalized to the X/XOD-derived control signal. Accordingly, a counter screen has been performed with cytochrome C or luminol without X/XO-generated superoxide.

### H_2_O_2_ measurement in cell-free system

The fluorescence of Amplex Red was measured in presence and absence of H_2_O_2_ (0.25 μM), reflecting NOX4 and NOX5 output. After addition of inhibitors and 50 μl Amplex Red reaction mixture containing 100 U•mL-1 HRP and 10 mM Amplex Red to each well, the plate was incubated at 37°C for 10 min. Thereafter, the plate was read at excitation (530-560 nm) and emission ~ 590 nm at 37°C and for 60 min. Data was calculated as the AUC over 60 min. and data were normalized to H_2_O_2_ output in absence of inhibitor.

### Cell permeability in HBMECs

For the passive dye diffusion assay, 2 × 10^4^ human brain microvascular endothelial cells (HBMECs) were grown to confluence on membranes of Transwell inserts (collagen-coated Transwell Pore Polyester Membrane Insert; pore size = 3.0 μm (Corning, The Netherlands) Fig. 5A). After 6hrs of ischemia, cells were treated with 0,1 μM ML171, 0,6 μM VAS2870, 1 μM GKT136901, 0,2 μM M13 or 0,01 μM ML090 during 24hrs of re-oxygenation. Thereafter, cell permeability was assessed using Evans Blue (Sigma-Aldrich, The Netherlands) (Fig. 5B). First, the medium was removed and previously warmed (37°C) PBS (1,5 ml) was added to the abluminal side of the insert. Permeability buffer (0,5 ml) containing 4% bovine serum albumin (Sigma-Aldrich, The Netherlands) and 0,67 mg/ml Evans blue dye in PBS was loaded on the luminal side of the insert followed by 15 min. incubation at 37°C. The concentration of Evans Blue in the abluminal chamber was measured by determining the absorbance of 200 μl buffer at 630 nm using a microplate reader.

### Statistical analysis

Data analysis was performed using Prism 6.0g software package (GraphPad Software, San Diego, USA). IC_50_ values were calculated with a non-linear regression analysis using an algorithm for sigmoidal dose–response with variable slopes. Results are expressed as means ± SEM. Statistical differences between means were analyzed by one-way ANOVA followed by Bonferroni correction for multiple comparisons. A value of P < 0.05 was considered statistically significant.

## Results

### Gene expression of NOX isoforms

To check the expression of the different NOX isoforms, we analysed NOX1, 2, 4 and 5 mRNA expression in control vector-transfected HEK293 cells, HEK293 cells transfected with NOX1, 2, 4 and 5 and HL-60 cells. NOX isoforms mRNA expression was measured by RT-qPCR and mRNA levels were normalized to the expression of β-actin.

As reported previously [26, 32], NOX1 was expressed in NOX1-transfected HEK293 cells (Fig. 1A), NOX2 in HL-60 (Fig. 1B), NOX4 in NOX4-transfected HEK293 (Fig. 1C) and NOX5 in NOX5-transfected HEK293 (Fig. 1D) and were below detection limit (defined as 100 times lower than in the respective NOX overexpressing cells) in all other cell preparations. Notably, the expression of NOX isoforms did not change after treatment with the respective assay conditions i.e. luminol for NOX1, cytochrome C for NOX2 and Amplex Red for NOX4 and NOX5 (Fig. 1A–1D).

**Figure 1:**
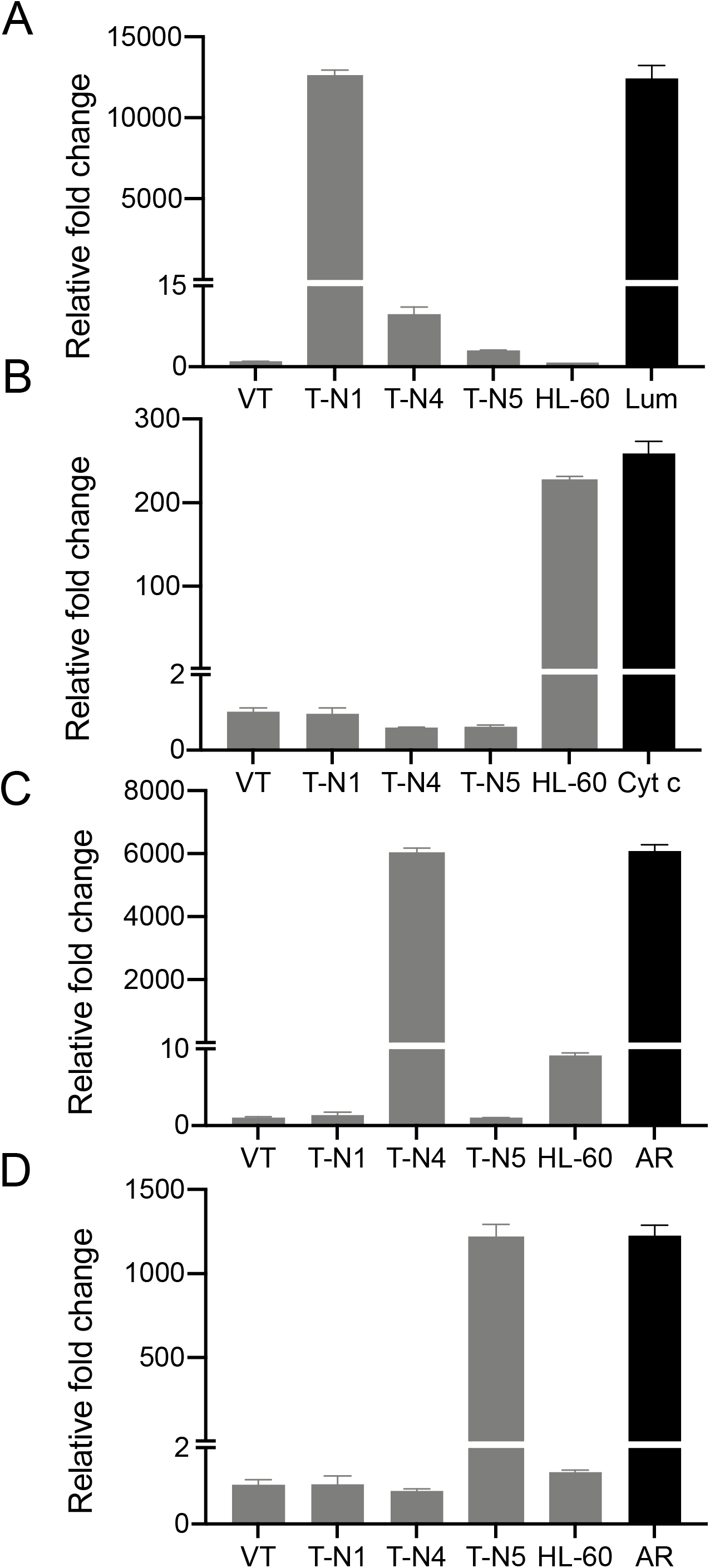
RT-qPCR of NOX isoforms mRNA expression in HEK293 and HL-60 cells. NOX isoforms mRNA levels were normalized to the expression of β-actin. To check whether assay reagents interfere with NOXs gene expression, HEK293 cells transfected with NOX1 (T-N1) were treated with luminol (Lum), HL-60 cells were treated with cytochrome C (Cyt c) and HEK293 cells transfected with NOX4 (T-N4) or NOX5 (T-N5) were treated with Amplex Red (AR). A) mRNA levels of NOX1. B) mRNA levels of NOX2. C) mRNA levels of NOX4. D) mRNA levels of NOX5. Data are presented as the mean±SEM of three experiments. VT, vector-transfected HEK293.

### NOX pharmacological isoform selectivity

To characterize the NOX isoform selectivity of the current second-generation NOX inhibitors we determined concentration-dependency and efficacy of GKT136901, Ml171, VAS2870, M13 and ML090 on NOX1, 2, 4, and 5 (Fig. 2A). Our preference was for cell (lines) expressing a specific NOX isoform in a highly selective manner and ideally physiologically. The latter is only known for NOX2 and HL-60 cells [33, 34]. For NOX1, NOX4 and NOX5, there is, to our knowledge, no such native human cell line. Therefore, for these three isoforms we used transiently transfected HEK-293 cells, using previously validated methodology [27].

Specifically, NOX2 expressing HL-60 cells or HEK293 cells transfected with NOX1, NOX4 or NOX5 were incubated in presence of increasing concentrations of each compound. ROS generation was subsequently induced by PMA to stimulate NOX1, NOX2 and NOX5, which was assayed in presence of ionomycin. Thereafter, ROS production was assayed using a panel of cellular assays with structurally unrelated probes; *i.e.* Amplex red, luminol or cytochrome C (Figure 2B). Amplex Red was used to quantify extracellular H_2_O_2_ [35], while luminol-based chemiluminescence and cytochrome C-reduction were used to quantify superoxide generation. As controls, medium, vector-transfected HEK293 cells and non-stimulated HL-60 cells were used (Fig. S1),

**Figure 2:**
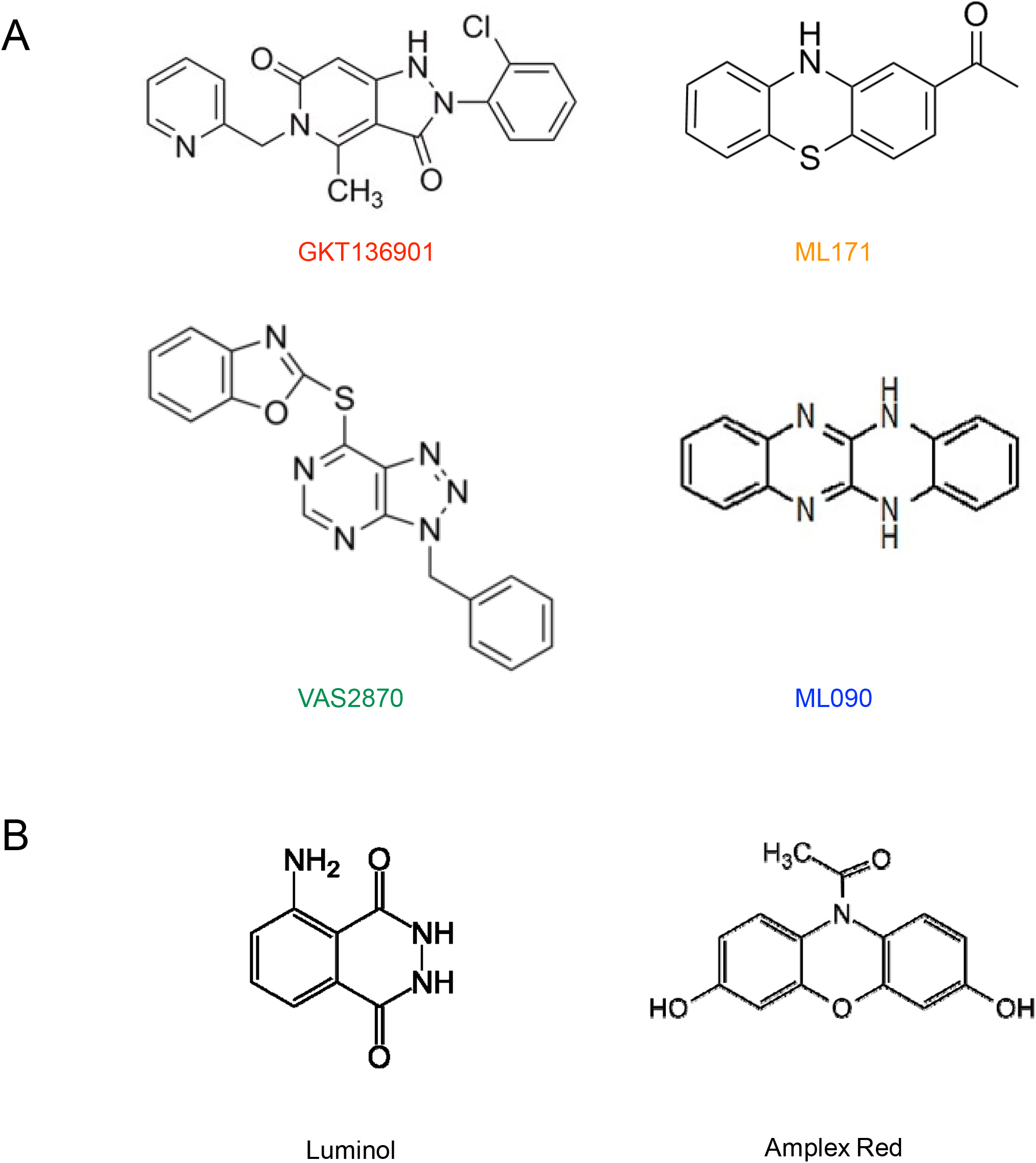
Chemical structures. Chemical structure of NOX inhibitors (A) and probes (B) used in the study.

NOX isoform concentration-response curves for GKT136901, Ml171, VAS2870, M13 and ML090 were constructed (Fig. 3; see table 1 for IC_50_ values). GKT136901 (Fig. 3A) showed selectivity for NOX1 over NOX4 and NOX5 inhibition while NOX2 inhibition was not observed. The same holds true for ML171 (Fig. 3B) which is more selective for NOX1 compared to NOX4 and NOX5. VAS2870 (Fig. 3C) displayed NOX2 over NOX1 and NOX4 selective inhibition and also slightly inhibited NOX5. The GlucoxBiotech compound M13 (Fig. 3D) showed almost selective NOX4 inhibition but NOX2 (with low E_max_) and NOX1 (at high concentrations) inhibition was observed as well. Finally, ML090 (Fig. 3E) inhibited NOX5, NOX1 and NOX4 with comparable IC_50_ values but enhanced E_max_ for NOX5 inhibition. These data suggest that NOX inhibitors indeed display differential isoform targeting, albeit the single compound selectivity is yet not sufficient. Therefore, combined analysis of a NOX inhibitor panel with sufficient differential potencies and selectivities is suggested.

**Figure 3:**
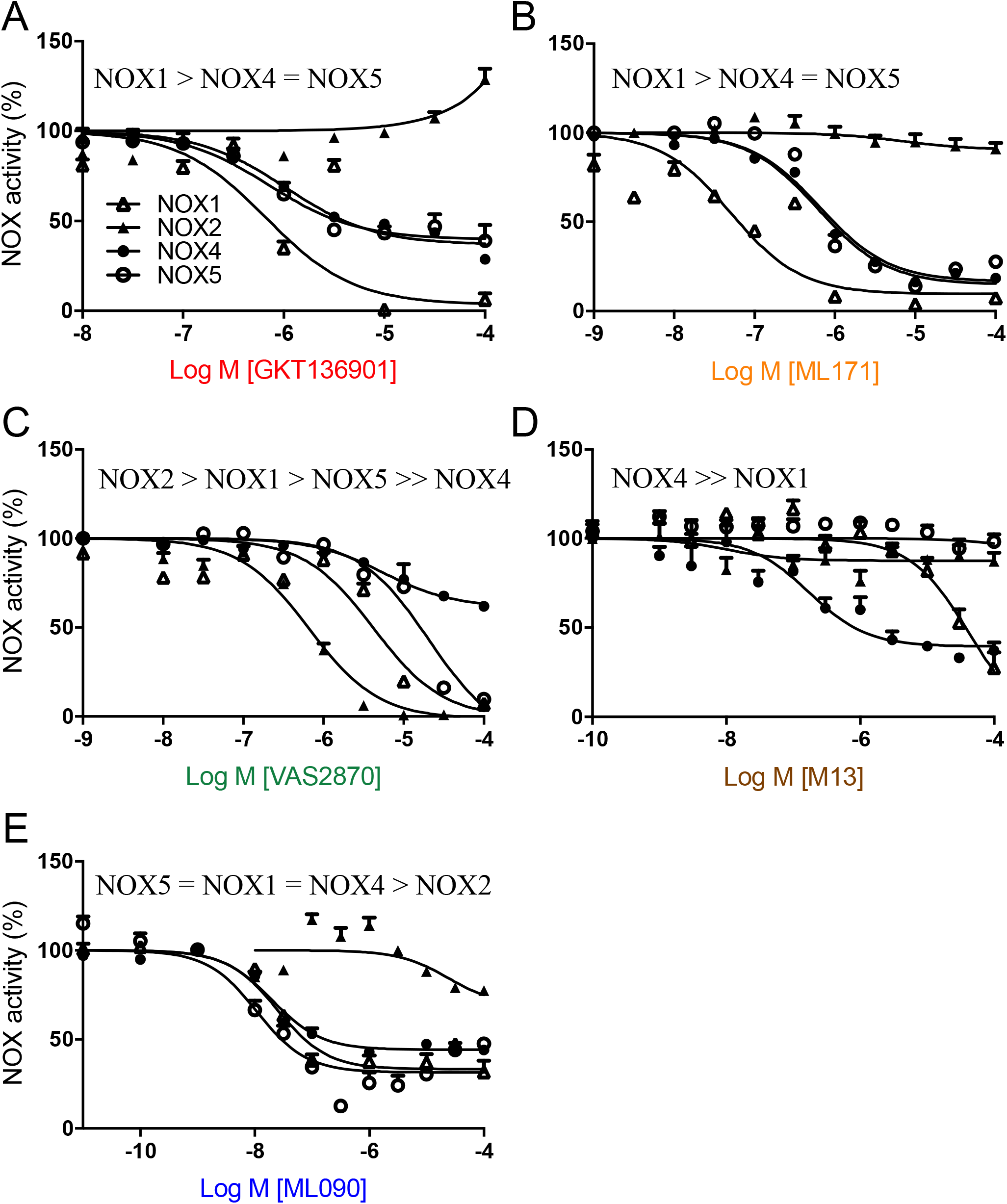
NOX inhibitors display isoform selectivity. ROS production was assessed in whole cells using assays measuring superoxide production by NOX1 (luminol assay) or NOX2 (cytochrome C assay) and H_2_O_2_ production by NOX4 or NOX5 (Amplex Red assay). As illustrated in the figure (A-E), GKT136901 (A), ML171 (B), VAS2870 (C), M13 (D) or ML090 (E) inhibited ROS production with NOX isoform selective IC_50_. Data are presented as the mean±SEM of three experiments.

**Table 1:** IC_50_ (Log M) and E_max_ (% of control) values for NOX isoform inhibition.

|  |  | NOX1 | NOX2 | NOX4 | NOX5 |
| --- | --- | --- | --- | --- | --- |
| GKT136901 | IC<sub>50</sub> | -6.17 | > -3 | -5.97 | -6.14 |
|  | E<sub>max</sub> | 99 | NA | 71 | 61 |
| ML171 | IC<sub>50</sub> | -7.3 | -5.16 | -6.21 | -6.18 |
|  | E<sub>max</sub> | 96 | 09 | 84 | 86 |
| VAS2970 | IC<sub>50</sub> | -5.37 | -6.18 | -5.25 | -4.69 |
|  | E<sub>max</sub> | 93 | 99 | 38 | 90 |
| M13 | IC<sub>50</sub> | -4.39 | -7.98 | -6.76 | -3.9 |
|  | E<sub>max</sub> | 73 | 24 | 67 | 05 |
| ML090 | IC<sub>50</sub> | -7.6 | -4.62 | -7.69 | -7.96 |
|  | E<sub>max</sub> | 68 | 23 | 57 | 81 |

### Non-specific anti-oxidant and assay artefacts

To screen for possible assay interference between GKT136901, ML171, VAS2870, M13 or ML090 and the NOX1 assay, a cell-free system was performed in which each inhibitor was screened in presence of luminol, 1mU/ml xanthine oxidase (XO) and 1mg xanthine (X) generating superoxide. At concentrations 1μM GKT136901 did not show interference with either the molecular probe or with the X/XO system (Fig. 4A). Similar findings were obtained with 0.1 μM ML171, 10 μM VAS2870 and 30 nM ML090 (Fig. 4A). In contrast, 1 μM M13 enhanced chemiluminescence (Fig. 4A). To identify direct interactions between assay components and GKT136901, ML171, VAS2870, M13 or ML090 a cell-free counter screen was performed without X/XO-generated ROS. Indeed, ML171 and VAS2870 showed reduction of the luminol-based signal suggesting direct interference with luminol-based chemiluminescence (Fig. 4B). In contrast, in presence of ML090 the signal was enhanced (Fig. 4B).

**Figure 4:**
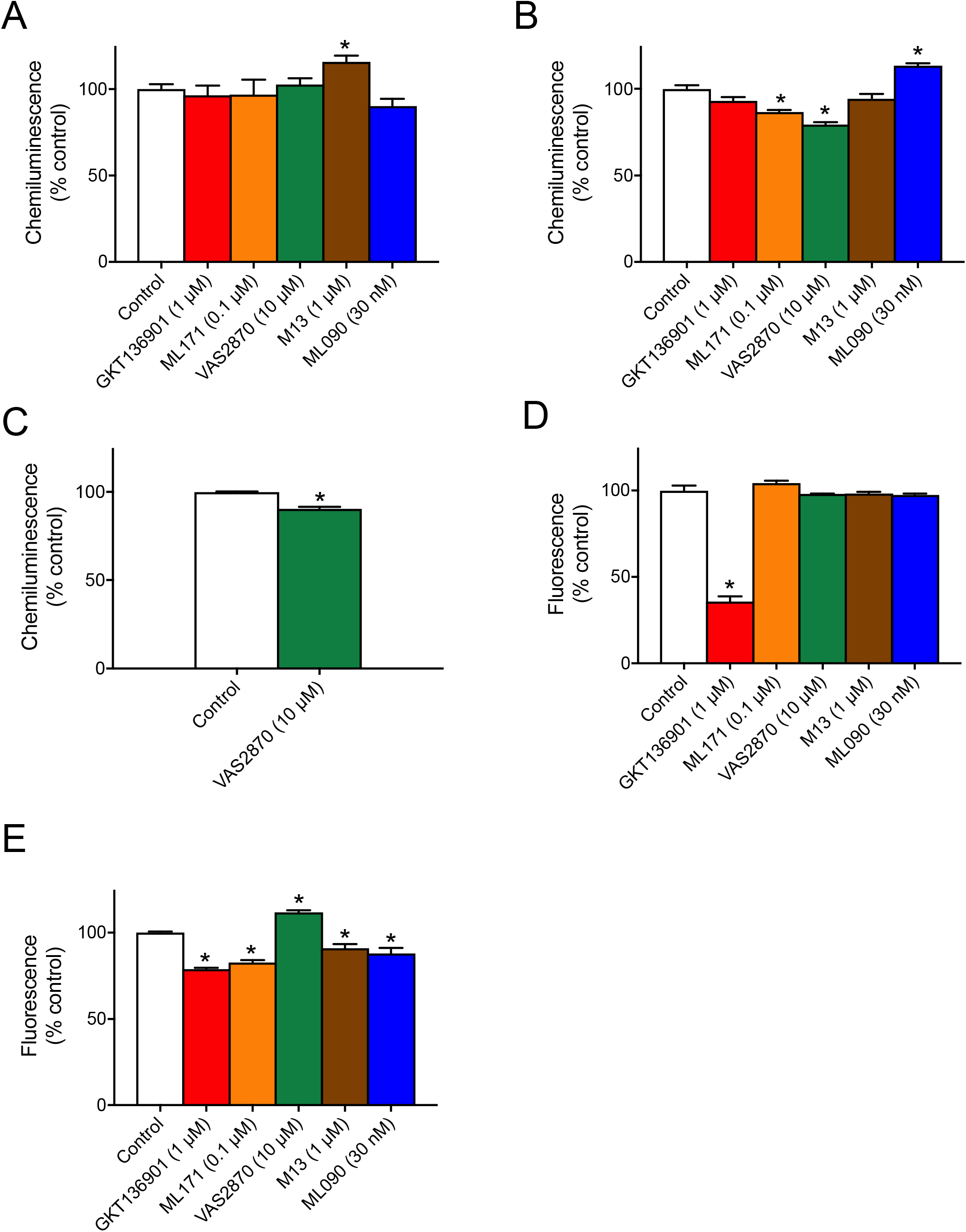
NOX inhibitors display interference with ROS assays. A/B) To study possible interference of GKT136901, ML171, VAS2870, M13 or ML090 with luminol-based measurements cell-free luminol assays were used. In these assays, ROS production generated by X/XO was enhanced by M13 (A) but not by GKT136901, ML171, VAS2870 or ML090. Moreover, chemiluminescence produced by luminol only was inhibited by ML171 and VAS2870, enhanced by ML090 and not affected by GKT136901 or M13, respectively (B). C) Possible assay interference by NOX2-specific VAS2870 was assessed by studying cytochrome C reduction in presence of X/XO-derived ROS in a cell-free system. Presence of VAS2870 slightly, but significantly reduced chemiluminescence. D/E) The possibility of interference between GKT136901, ML171, VAS2870, M13 or ML090 and Amplex Red-based assays was studied using cell-free Amplex Red assays. In these assays, Fluorescence generated by H_2_O_2_ was significantly inhibited by GKT136901 (D) but not by ML171, VAS2870, M13 or ML090 (D). Moreover, fluorescence produced by Amplex Red only was inhibited by GKT136901, ML171, M13 and ML090 while it was enhanced by VAS2870 (E). Data are presented as the mean±SEM of three experiments. * P < 0.05.

To study non-specific antioxidant effects of VAS2870, the effect of 10 μM VAS2870 was studied in a cell-free system in presence of cytochrome C, and X/XO-derived ROS. In these assays, VAS2870 showed significant antioxidant effects (Fig. 4C).

To study whether the effects of GKT136901, ML171, VAS2870, M13 and ML090 on NOX4 and NOX5 are specific, a cell-free assay was performed with NOX4 or NOX5 effective concentrations of each inhibitor in presence of 0.25μM H2O2 in an Amplex Red assay. ML171, VAS2870, M13 and ML090 did not directly affect Amplex Red-based detection of H_2_O_2_ (Fig. 4D). However, GKT136901 reduced Amplex Red-based H_2_O_2_ signals suggesting assay interference. Hence, the assay was repeated without H_2_O_2_ to study potential direct assay component interference. Indeed, GKT136901, ML171, M13 and ML090 reduced the signal as compared to the control (Fig. 4E) whereas in presence of VAS2870 the signal was enhanced (Fig. 4E).

### Target validation of NOX in ischemia-induced hyperpermeability model

As predicted from the inhibitor screen, M13, GKT136901 or ML171 protect against ischemia-induced hyperpermeability in HBMEC. Our data suggested that a NOX inhibitor panel may be used for target validation of selective NOX isoforms. We thus tested our NOX inhibitor panel in an *in vitro* human model of ischemia-induced hyperpermeability (Fig. 5A) where NOX4 is involved in subacute hypoxia-induced increases in cell permeability whereas NOX1, NOX2 do not [31] and NOX5 only acutely [20]. Primary HBMEC cultures were subjected to 6h of hypoxia followed by 24h of re-oxygenation (Fig. 5B) in presence or absence of ML171 (0.1 μM; mainly targeting NOX1), VAS203 (0.6 μM; mainly targeting NOX2), GKT136901 (1 μM; mainly targeting NOX4), M13 (0.2 μM; mainly targeting NOX4) or ML090 (0.01 μM; mainly targeting NOX5). Hypoxia increased cell permeability after 24hrs of re-oxygenation. GKT136901, and ML171 treatment prevented this detrimental effect (Fig. 5C), suggesting protection against hyperpermeability via NOX4 inhibition while, as expected given their respective IC_50_ values, VAS2870 or ML090 treatment showed no effect (Fig. 5C). These data provide a proof-of-concept for pharmacological target validation using NOX inhibitor panels.

**Figure 5:**
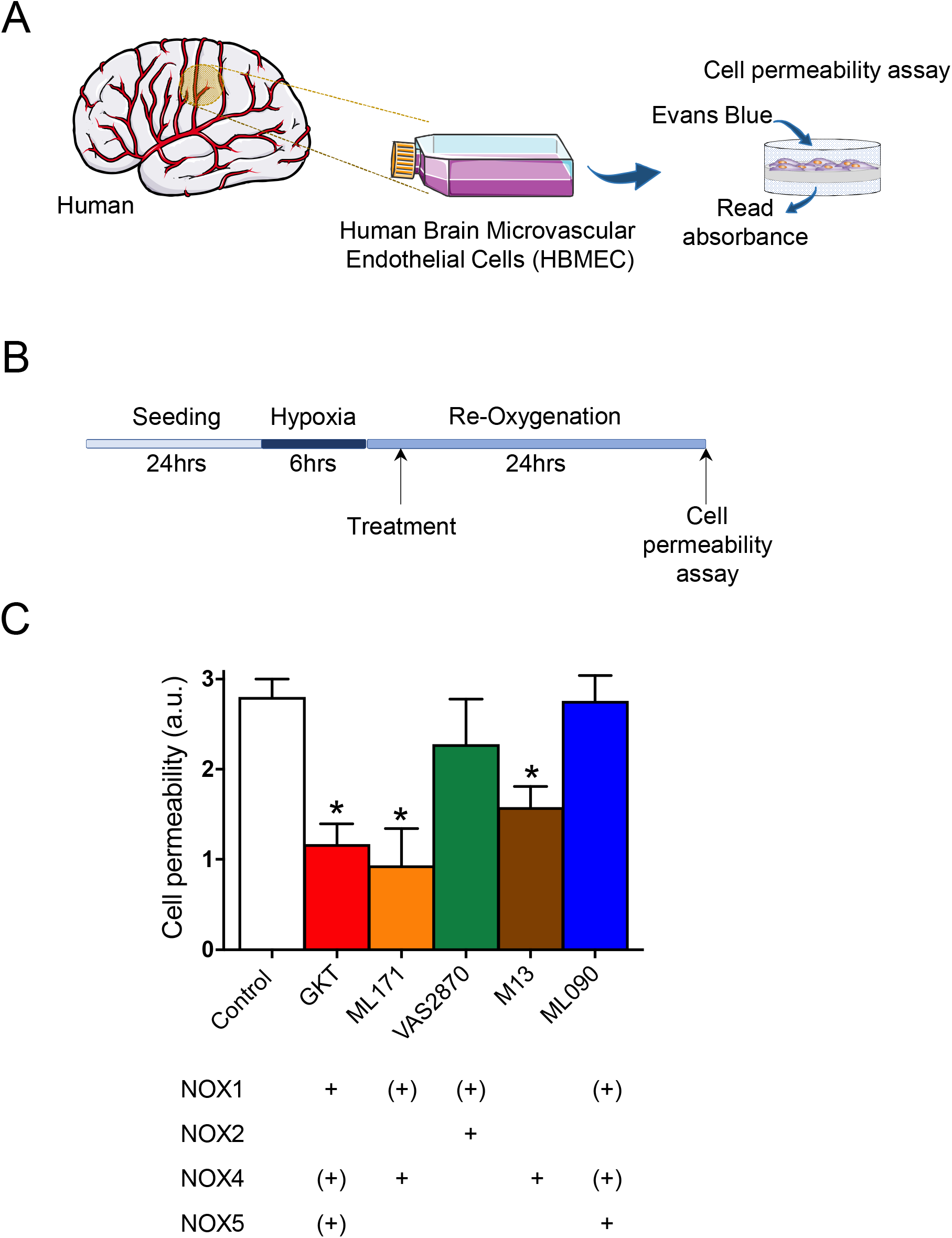
GKT136901, GKT136901 and M13 inhibit ischemia-induced hyperpermeability. A/B) HCMECs were subjected to 6hrs hypoxia followed by 24hrs re-oxygenation during which inhibitors were present. Cell permeability was subsequently assessed by measuring Evans Blue (EB) fluorescence. C) Hypoxia caused hyperpermeability (open bar) which was significantly reduced in cells treated with ML171 (0.1 μM), GKT136901 (0.1 μM) or M13 (0.2 μM) while VAS2870 (0.6 μM) or ML090 (0.01 μM) treatment had no effect. Data are presented as the mean±SEM of 4-6 experiments; * P < 0.05.

## Discussion

Our results show that NOX-specific and isoform-selective pharmacological NOX inhibition is in principle achievable. Individual compound specificity is yet insufficient. None of the selected small molecule inhibitors were sufficiently isoform selective but inhibited several NOX isoforms with different ranking potencies. In addition, assay interferences, some of them previously not noted, needed to be taken into consideration. Nevertheless, using a panel of NOX inhibitors pharmacological validation of a specific NOX isoform was possible and examplified in a NOX4-dependent model.

With respect to the individual NOX inhibitor compounds in this study, VAS2870 showed beneficial effects in several preclinical disease models such as stroke [31], Alzheimer’s disease [36], thrombosis [37] and pulmonary hypertension [38]. It has been designated a pan-NOX inhibitor [39] as inhibition of NOX1 [26, 40], NOX2 [26, 40, 41], NOX4 [26, 31] and NOX5 [26] activity has been observed; IC_50_ values however have been only published for NOX2 [41, 42]. Based on the here reported IC_50_ for all tested NOX isoforms, VAS2870 is relatively NOX2 selective (IC_50_ ~ 0.7μM), followed by NOX1 and 4 in the lower and NOX5 in the higher micromolar range. VAS2870 showed interference with luminol and Amplex Red assays and reduced XO-generated ROS indicating antioxidant properties as reported previously [25]. Interference with XO-generated ROS was 9.6% (Fig. 4C); the same concentration, however, resulted in a 91.5% decrease of the NOX2-dependent signal (Fig. 3C).

M13, a previously unreported compound from Glucox Biotech, was identified as a first-in-class relative NOX4 selective inhibitor as it was 200-times more potent in inhibiting NOX4 (IC_50_ ~ 0.01 μM) versus NOX1 (IC_50_ ~ 0.2 μM) and showed negligible NOX2 inhibition and almost no effect on NOX5. Although M13 interfered with the Amplex Red assay by 9% (Fig. 4E), this was considerably less than the 40% reduction of the NOX4-dependent signal (Fig. 3D).

ML090 was previously described as NOX1 selective [43]. In our hands, IC_50_ values for NOX1, 4 and 5 were quite similar suggesting ML090 to be a rather pan NOX inhibitor. Two interferences were observed. The 36.8% inhibition of the NOX1-dependent signal (Fig. 3D) may represent a slight underestimation because ML090 per se surprisingly increased the luminol signal by 13.4% (Fig. 4B). ML090 also interfered with the Amplex Red signal, yet lowering it by 11.9% (Fig. 4E); the NOX5 dependent signal, however, was reduced by 46.8 % (Fig. 3E). Therapeutically, ML090 protects from vascular dysfunction in a rabbit model, an effect that has been attributed to NOX1 inhibition [44]. According to our here presented data, however, an effect involving NOX4 and, in particular, NOX5 cannot be excluded as rabbits, unlike mice and rats, do express this isoform.

Compared to all compounds, GKT136901, a chemical analogue and pharmacological sibling of the clinically most advanced NOX inhibitor, GKT831, and claimed to be NOX1 and NOX4 specific, stood out in a problematic manner. GKT136901interfered both with the Amplex Red assay, for NOX4 and 5, and the luminol assay, for NOX1, and, on top of this, directly scavenged ROS. This latter antioxidant effect has been observed also by others using an alternative H_2_O_2_ assay [45] or by scavenging peroxynitrite [28]. These characteristics thus appear to limit interpretation of results using GKT136901 as a NOX4/5 inhibitor and may cast doubt on the specificity with respect to NOX of the entire GKT compound family in general-.

Similarly, the phenothiazine derivative, ML171, which had been published as NOX1 specific [46] and beneficial effects in models of hypertension [47], diabetes [48] and cancer [49], showed assay interferences with both Amplex Red and the luminol assay in line with a previous observation [50] that phenothiazines are peroxidase substrates [51]. Nevertheless, the NOX1-dependent chemiluminescence signal was inhibited to a larger extend than the assay interference.

Thus, results from the detection of NOX-derived ROS need careful interpretation and several controls as most of probes and inhibitors are prone to artifacts (see als review in [52]). Facing these individual compound limitations, we moved to a NOX inhibitor panel used at IC_50_ concentrations for to test whether they would allow pharmacological target validation of a specific NOX isoform. Indeed, using this approach our observation that GKT136901, ML171 and M13 were neuroprotective, but ML090 and VAS2870 were not, suggested NOX4 as responsible isoform, which had previously been established genetically [15]. Thus, the method is inhibitor panel approach in principle feasible for pharmacological target validation.

In summary, all tested NOX inhibitor compounds displayed different isoform selectivity profiles suggesting differential chemical targeting and a proof-of-principle that NOX isoform selective small molecule inhibition will become possible. This should stimulate focused library synthesis and structure-activity programs for further lead optimization to obtain truly isoform selective small molecules for single compound target validation and potential for clinical development. For now, we provide an immediately applicable inhibitor panel approach that allows target validation of NOXs under conditions where gene knock-out or knock-in are not feasible or, because of compensation mechanism, not desirable.

## Acknowledgements

We wish to thank Dr. Per Wikström for providing M13, Dr. V. Jaquet for experimental advice and Dr. Merlijn J. Meens for reading the manuscript and helpful discussions. Technical assistance by P. Lijnen and J. Bost is gratefully acknowledged. Financial support (to HHHWS) by the ERC (AdG RadMed and PoC SAVEBRAIN) and the Horizon 2020 programme (REPO-TRIAL) is gratefully acknowledged.

## Author Contributions

H.H.H.W.S., V.T.D, S.A and M.H.E designed research; V.T.D., S.A, M.H.E, P.L, A.I.C, M.P.P, C.N and U.K performed research; V.T.D, S.A, A.I.C, M.H.E and H.H.H.W.S. analyzed and interpreted data; V.T.D and M.H.E contributed in writing and revised the final manuscript and figures; M.H.E, V.T.D. and H.H.H.W.S. wrote and edited figures and manuscript.

---

**Figure S1.**
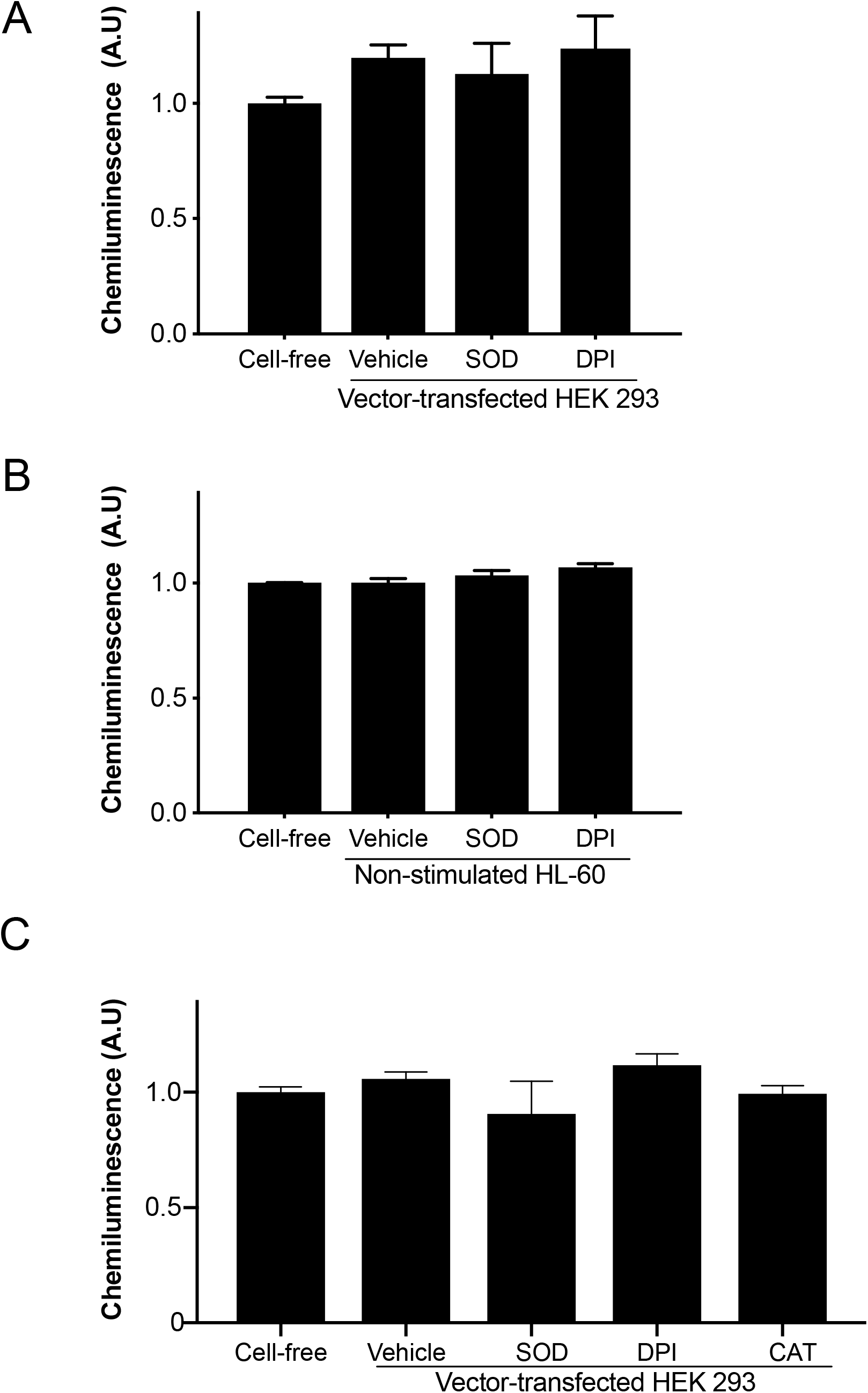

## References

1. Dotan, Y. et al. (2009) No evidence supports vitamin E indiscriminate supplementation. BioFactors (Oxford, England) 35 (6), 469–473.

2. Miller, E.R. et al. (2005) Meta-analysis: high-dosage vitamin E supplementation may increase all-cause mortality. Annals of internal medicine 142 (1), 37–46.

3. Bjelakovic, G. et al. (2007) Mortality in randomized trials of antioxidant supplements for primary and secondary prevention: systematic review and meta-analysis. JAMA 297 (8), 842–857.

4. Investigators, H.O.P.E.S. et al. (2000) Vitamin E supplementation and cardiovascular events in high-risk patients. The New England journal of medicine 342 (3), 154–160.

5. Wingler, K. et al. (2011) NOX1, 2, 4, 5: counting out oxidative stress. British journal of pharmacology 164 (3), 866–883.

6. Bedard, K. and Krause, K.-H. (2007) The NOX family of ROS-generating NADPH oxidases: physiology and pathophysiology. Physiological reviews 87 (1), 245–313.

7. Ray, P.D. et al. (2012) Reactive oxygen species (ROS) homeostasis and redox regulation in cellular signaling. Cellular signalling 24 (5), 981–990.

8. Finkel, T. (2011) Signal transduction by reactive oxygen species. The Journal of cell biology 194 (1), 7–15.

9. Dao, V.T. et al. (2015) Pharmacology and Clinical Drug Candidates in Redox Medicine. Antioxidants & redox signaling 23 (14), 1113–1129.

10. Elbatreek, M.H. et al. (2019) Reactive Oxygen Comes of Age: Mechanism-Based Therapy of Diabetic End-Organ Damage. Trends in endocrinology and metabolism: TEM 30 (5), 312–327.

11. Casas, A.I. et al. (2015) Reactive Oxygen-Related Diseases: Therapeutic Targets and Emerging Clinical Indications. Antioxidants & redox signaling 23 (14), 1171–1185.

12. Gray, S.P. et al. (2013) NADPH oxidase 1 plays a key role in diabetes mellitus-accelerated atherosclerosis. Circulation 127 (18), 1888–1902.

13. Wilkinson-Berka, J.L. et al. (2013) NADPH Oxidase, NOX1, Mediates Vascular Injury in Ischemic Retinopathy. Antioxidants & redox signaling.

14. Sorce, S. and Krause, K.H. (2009) NOX enzymes in the central nervous system: from signaling to disease. Antioxidants & redox signaling 11 (10), 2481–2504.

15. Casas, A.I. et al. (2017) NOX4-dependent neuronal autotoxicity and BBB breakdown explain the superior sensitivity of the brain to ischemic damage. Proceedings of the National Academy of Sciences of the United States of America 114 (46), 12315–12320.

16. Jha, J.C. et al. (2014) Genetic Targeting or Pharmacologic Inhibition of NADPH Oxidase Nox4 Provides Renoprotection in Long-Term Diabetic Nephropathy. Journal of the American Society of Nephrology: JASN.

17. Geis, C. et al. (2017) NOX4 is an early initiator of neuropathic pain. Experimental neurology 288, 94–103.

18. Gray, S.P. et al. (2016) Reactive Oxygen Species Can Provide Atheroprotection via NOX4-Dependent Inhibition of Inflammation and Vascular Remodeling. Arteriosclerosis, Thrombosis, and Vascular Biology 36 (2), 295–307.

19. Schürmann, C. et al. (2015) The NADPH oxidase Nox4 has anti-atherosclerotic functions. European heart journal 36 (48), 3447–3456.

20. Casas, A.I. et al. (2019) Calcium-dependent blood-brain barrier breakdown by NOX5 limits postreperfusion benefit in stroke. The Journal of clinical investigation 130 (4), 1772–1778.

21. Holterman, C.E. et al. (2013) Nephropathy and Elevated BP in Mice with Podocyte-Specific NADPH Oxidase 5 Expression. Journal of the American Society of Nephrology: JASN, ASN.2013040371.

22. Jha, J.C. et al. (2017) NADPH Oxidase Nox5 Accelerates Renal Injury in Diabetic Nephropathy. Diabetes 66 (10), 2691–2703.

23. Pandey, D. et al. (2012) Expression and functional significance of NADPH oxidase 5 (Nox5) and its splice variants in human blood vessels. American journal of physiology. Heart and circulatory physiology 302 (10), H1919–28.

24. Guzik, T.J. et al. (2008) Calcium-dependent NOX5 nicotinamide adenine dinucleotide phosphate oxidase contributes to vascular oxidative stress in human coronary artery disease. Journal of the American College of Cardiology 52 (22), 1803–1809.

25. Altenhöfer, S. et al. (2015) Evolution of NADPH Oxidase Inhibitors: Selectivity and Mechanisms for Target Engagement. Antioxidants & redox signaling 23 (5), 406–427.

26. Altenhöfer, S. et al. (2012) The NOX toolbox: validating the role of NADPH oxidases in physiology and disease. Cell Mol Life Sci 69 (14), 2327–2343.

27. Heumüller, S. et al. (2008) Apocynin is not an inhibitor of vascular NADPH oxidases but an antioxidant. Hypertension 51 (2), 211–217.

28. Schildknecht, S. et al. (2014) The NOX1/4 inhibitor GKT136901 as selective and direct scavenger of peroxynitrite. Current medicinal chemistry 21 (3), 365–376.

29. Zielonka, J. et al. (2014) High-throughput assays for superoxide and hydrogen peroxide: design of a screening workflow to identify inhibitors of NADPH oxidases. The Journal of biological chemistry 289 (23), 16176–16189.

30. Zielonka, J. et al. (2013) On the use of L-012, a luminol-based chemiluminescent probe, for detecting superoxide and identifying inhibitors of NADPH oxidase: a reevaluation. Free radical biology & medicine 65, 1310–1314.

31. Kleinschnitz, C. et al. (2010) Post-stroke inhibition of induced NADPH oxidase type 4 prevents oxidative stress and neurodegeneration. PLoS biology 8 (9).

32. Wind, S. et al. (2010) Comparative pharmacology of chemically distinct NADPH oxidase inhibitors. British journal of pharmacology 161 (4), 885–898.

33. Teufelhofer, O. et al. (2003) Promyelocytic HL60 cells express NADPH oxidase and are excellent targets in a rapid spectrophotometric microplate assay for extracellular superoxide. Toxicol Sci 76 (2), 376–83.

34. Hirano, K. et al. (2015) Discovery of GSK2795039, a Novel Small Molecule NADPH Oxidase 2 Inhibitor. Antioxid Redox Signal 23 (5), 358–74.

35. Maghzal, G.J. et al. (2012) Detection of reactive oxygen species derived from the family of NOX NADPH oxidases. Free radical biology & medicine 53 (10), 1903–1918.

36. Abubaker, A.A. et al. (2019) Amyloid Peptide beta1-42 Induces Integrin alphaIIbbeta3 Activation, Platelet Adhesion, and Thrombus Formation in a NADPH Oxidase-Dependent Manner. Oxid Med Cell Longev 2019, 1050476.

37. Avdonin, P.V. et al. (2019) VAS2870 Inhibits Histamine-Induced Calcium Signaling and vWF Secretion in Human Umbilical Vein Endothelial Cells. Cells 8 (2).

38. Li, T. et al. (2019) Magnesium lithospermate B prevents phenotypic transformation of pulmonary arteries in rats with hypoxic pulmonary hypertension through suppression of NADPH oxidase. Eur J Pharmacol 847, 32–41.

39. Wingler, K. et al. (2012) VAS2870 is a pan-NADPH oxidase inhibitor. Cell Mol Life Sci 69 (18), 3159–3160.

40. Wind, S. et al. (2010) Oxidative stress and endothelial dysfunction in aortas of aged spontaneously hypertensive rats by NOX1/2 is reversed by NADPH oxidase inhibition. Hypertension 56 (3), 490–497.

41. Gatto, G.J. et al. (2013) NADPH oxidase-dependent and - independent mechanisms of reported inhibitors of reactive oxygen generation. Journal of enzyme inhibition and medicinal chemistry 28 (1), 95–104.

42. ten Freyhaus, H. et al. (2006) Novel Nox inhibitor VAS2870 attenuates PDGF-dependent smooth muscle cell chemotaxis, but not proliferation. Cardiovascular research 71 (2), 331–341.

43. Brown, S.J. et al. (2010) Probe Report for NOX1 Inhibitors.

44. Smith, R.M. et al. (2015) Role of Nox inhibitors plumbagin, ML090 and gp91ds-tat peptide on homocysteine thiolactone induced blood vessel dysfunction. Clin Exp Pharmacol Physiol 42 (8), 860–4.

45. Martyn, K.D. et al. (2006) Functional analysis of Nox4 reveals unique characteristics compared to other NADPH oxidases. Cellular signalling 18 (1), 69–82.

46. Gianni, D. et al. (2010) A novel and specific NADPH oxidase-1 (Nox1) small-molecule inhibitor blocks the formation of functional invadopodia in human colon cancer cells. ACS chemical biology 5 (10), 981–993.

47. Harvey, A.P. et al. (2017) Vascular dysfunction and fibrosis in stroke-prone spontaneously hypertensive rats: The aldosterone-mineralocorticoid receptor-Nox1 axis. Life Sci 179, 110–119.

48. Weaver, J.R. et al. (2015) Inhibition of NADPH oxidase-1 preserves beta cell function. Diabetologia 58 (1), 113–21.

49. Liang, S. et al. (2019) NADPH Oxidase 1 in Liver Macrophages Promotes Inflammation and Tumor Development in Mice. Gastroenterology 156 (4), 1156–1172 e6.

50. Seredenina, T. et al. (2015) A subset of N-substituted phenothiazines inhibits NADPH oxidases. Free radical biology & medicine 86, 239–249.

51. Rogozhina, T.V. and Rogozhin, V.V. (2011) [Phenothiazines are slowly oxidizable substrates of horseradish peroxidase]. Biomeditsinskaia khimiia 57 (5), 544–553.

52. Maghzal, G.J. et al. (2012) Detection of reactive oxygen species derived from the family of NOX NADPH oxidases. Free Radic Biol Med 53 (10), 1903–18.

