## Supplemental information for "Isoform-selective NADPH oxidase inhibitor panel for pharmacological target validation"

**Isoform-specific NADPH oxidase inhibition for pharmacological target validation**

**Table s1: qPCR Assays**

| **Gene** | **Assay ID** |
| --- | --- |
| β-actin | Hs99999903_m1 |
| Nox1 | Hs00246589_m1 |
| Nox2 | Hs00166163_m1 |
| Nox4 | Hs01558199_m1 |
| Nox5 | Hs00225846_m1 |

**Figure S1:**

A

B

C

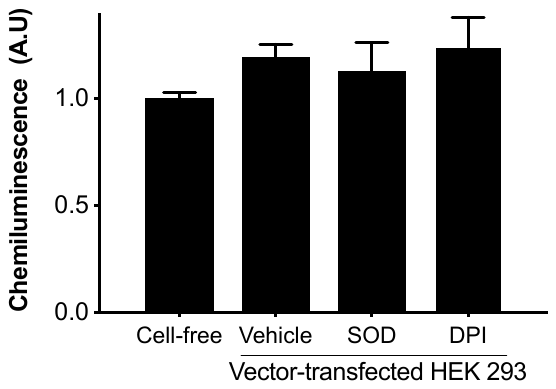

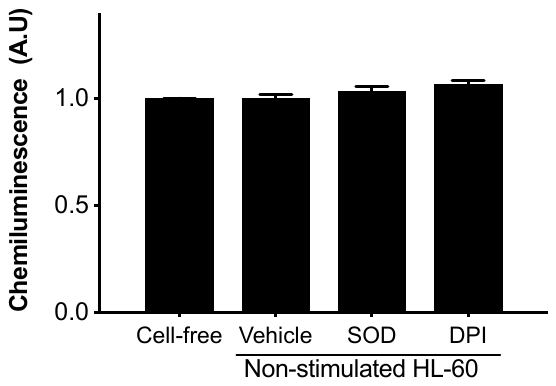

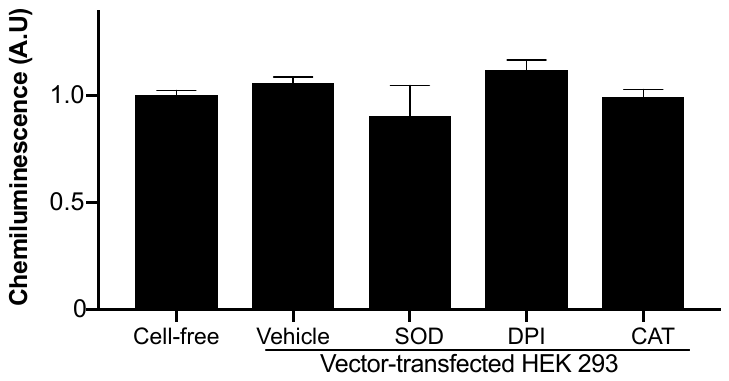

**Figure S1: Controls for ROS production.** Comparison of cell-free medium or control cells treated with the ROS inhibitors; SOD, catalase or diphenylene iodonium (DPI) in (A) Luminol assay, (B) Cytochrome C assay and (C) Amplex Red assay.
